# Global thermal tolerance of freshwater invertebrates and fish

**DOI:** 10.1101/2024.07.08.602306

**Authors:** Helena S. Bayat, Fengzhi He, Graciela M. Madariaga, Camilo Escobar-Sierra, Sebastian Prati, Jonathan F. Jupke, Kristin Peters, Xing Chen, Jurg W. Spaak, Alessandro Manfrin, Noel P.D. Juvigny-Khenafou, Ralf B. Schäfer

**Author notes:** Corresponding author: Helena S. Bayat.

## Abstract

Scientists have investigated the thermal tolerance of organisms for centuries, yet the field has not lost relevance as the environmental threats of thermal pollution and global change sharpen the need to understand the thermal vulnerability of organisms in landscapes increasingly subjected to multiple stressors. Freshwater fish and especially invertebrates are greatly underrepresented in recent large-scale compilations of thermal tolerance, despite the importance of freshwater habitats as a crucial resource and biodiversity havens. This inspired us to create a thermal tolerance database for these organisms that includes literature from 1900 until the present day sourced from five languages to counteract geographic bias, and 395 thermal tolerance tests conducted with additional stressors present. The database contains over 5000 records for over 800 species, including 452 invertebrates, providing a valuable resource to test hypotheses on thermal risks to freshwater organisms in present and future environments, and how these might change in multiple stressor scenarios.

## Background & Summary

Thermal limits of life have interested researchers since at least the 1700s^1–5^, with early comparisons of thermal tolerances between organisms published in the late 1800s^3^. Despite the large volume of research on the biological and ecological influence of temperature in the last two centuries, many questions regarding thermal tolerances remain unanswered^6–8^. For example, the relationship between oxygen and thermal tolerance is still contested^9,10^, and the influence of thermal tolerance on range limits not fully understood^11,12^. The evolution of thermal tolerance and to what extent it is conserved across the phylogeny of life has only been examined for selected groups^13–17^.

Additionally, thermal tolerance can be measured in different ways, which complicates large scale comparisons^18–22^. Two fundamental methodologies have emerged to measure thermal tolerance of a species in the last century. One method involves ramping the temperature until a specified endpoint is reached, known as the critical thermal method, coined by Cowles and Bogert in 1944^23^, though similar methods existed previously^24,25^. The metrics for upper and lower thermal tolerance are known as critical thermal maximum (CTmax) and critical thermal minimum (CTmin), respectively, which represent the upper and lower bounds of a thermal performance curve^26–28^. The most commonly measured endpoint for this method is the loss of equilibrium, which represents a temperature at which an organism can no longer escape conditions that would lead to death; however, an organism should be able to recover once removed from the test^19,29^. This method has gained popularity over time, as the test can be conducted quickly, without permanently harming the organisms, repeatedly on the same organism, and without large numbers of test organisms^29^. Modern mobile heating units allow this methodology to be conducted in the field^30,31^. The lethal thermal maximum (LTmax) or lethal thermal minimum (LTmin) metrics are reached when temperature is ramped until death occurs^32^. The second method is a static assay, wherein groups of organisms are kept at a fixed temperature for a fixed period of time. This method was formally named the incipient lethal method by Fry in 1947^33^, based on work by others in the two decades prior^34–38^. Percentage mortality is assessed as an endpoint for each group, and statistical methods such as probit analysis used to evaluate the temperature at which a certain percentage of the group is affected^37^. The LT50 metric is the temperature at which 50% of the organisms perish, analogous to the LC50 of ecotoxicological assays for chemicals^33,39^. As with LC50s, LT50s are commonly measured at 24, 48, or 96 hours^18^. The incipient upper or lower lethal temperature (IULT and ILLT, respectively) is a metric distinguished from the LT50 by the longer time course of the test – the IULT represents the temperature that 50% of a population would survive indefinitely^34^. This method has decreased in relative popularity over time, due to the large amount of resources needed^29^. A few studies have compared these two metrics on a theoretical^40,41^ and empirical^42^ level, but with limited sample sizes and no comparisons for fish to date.

With the advance of climate change, large collections of thermal limits have been compiled to assess the vulnerability of organisms to warming on a global scale and answer fundamental questions on the determinants of thermal tolerance^43–48^. Freshwater taxa are not well-represented within these collections, with the exemption of amphibians^43,48–50^. Freshwaters are critical habitats, both for ecological reasons and as a vital resource^51^. For example, freshwater ecosystems provide high quality subsidies to riparian and terrestrial ecosystems^52^. In addition, these systems provide services such as drinking water, water purification and contribute to climate regulation. Freshwater ecosystems are directly threatened by global warming patterns on a large scale, and locally by thermal pollution from industrial effluents ^53–55^. Moreover, climate change may also indirectly increase water temperatures. For example, flow intermittence induced by climatic changes may indirectly result in a strongly increased water temperatures^6,56^. Other stressors, such as hydromorphological changes and removal of riparian vegetation may also be associated with increases in water temperature^6,57,58^. To evaluate the effects of increasing temperatures requires information on their tolerance. Hence, we established a database focusing on the thermal tolerance of freshwater taxa, which allows for intra- and interspecific comparisons among global freshwater assemblages, assessment of thermal vulnerability under warming scenarios, and prediction of future range shifts.

The database was developed following current recommendations of best practice^59–61^. In detail, in comparison to previous studies we aimed to reduce geographic bias, provide multiple records per species (where possible, to allow assessment of intraspecific variation), and measures of dispersion, to address gaps also recently described by other authors^7,50^. We have included thermal tolerance metrics determined from the two main methodologies described above, including those conducted on the same species within the same study^42,62^, which facilitates a comparison of how the metrics are related across taxonomic groups. Our database focuses on freshwater fish and invertebrates, with records collated from previous work and expanded with new literature searches, including in relevant non-English languages. Including major non-English world languages in ecological literature searches decreases geographical bias, which is critical for extrapolating results across the diverse regions of the earth^63,64^. The current uncertainty induced by geographical bias in physiological data overrides the uncertainty in future climate predictions^65^, and non-English literature can help counteract this bias^66– 68^. Additionally, the publication rate of non-English studies on biodiversity is increasing in many countries^69^, thus the relevance of non-English literature may increase.

Thermal tolerance can also change with the addition of other stressors^49,70^, which affect its ecological relevance in a world where many organisms are increasingly subjected to multiple stressors^71,72^. We include thermal tolerance tests in the presence of additional stressors, with information on the type and level of the stressor(s). This facilitates future use of the data in untangling complex interactions between multiple stressors and consequential effects on organisms in stressful environments.

## Methods

### Inclusion criteria

The database includes studies that fulfill three primary criteria: 1) report at least one thermal tolerance measure for an organism, 2) have mortality or a sublethal indication of imminent mortality (such as loss of equilibrium, loss of righting response, or onset of spasms) as an endpoint, and 3) test organism(s) are fish or invertebrates residing in fresh or brackish water habitats for at least part of their life cycle (**Figure 1**). Studies were obtained by harmonizing existing databases, which automatically fulfilled the first two criteria and subsequently were filtered for relevant taxa. Additional studies were obtained through literature searches, wherein candidate studies were screened by title and abstract before data extraction. Studies that contained incomprehensible or missing methodology for thermal tolerance tests were excluded; minor methodological inconsistencies were noted during data extraction. The thermal tolerance metrics considered for inclusion needed to represent the result of either a dynamic or static assay, and either contain enough methodological information to deduce which kind of assay was conducted or cite the recognized metric name (CTmax, CTmin, LTmax, LTmin, LT50, IULT, ILLT) in accordance with literature describing methodology (notably key papers such as Cowles and Bogert 1944, Becker and Genoway 1979, Lutterschmidt and Hutchinson 1997, Beitinger and Bennett 2000)^18,19,23,29^.

**Figure 1.**
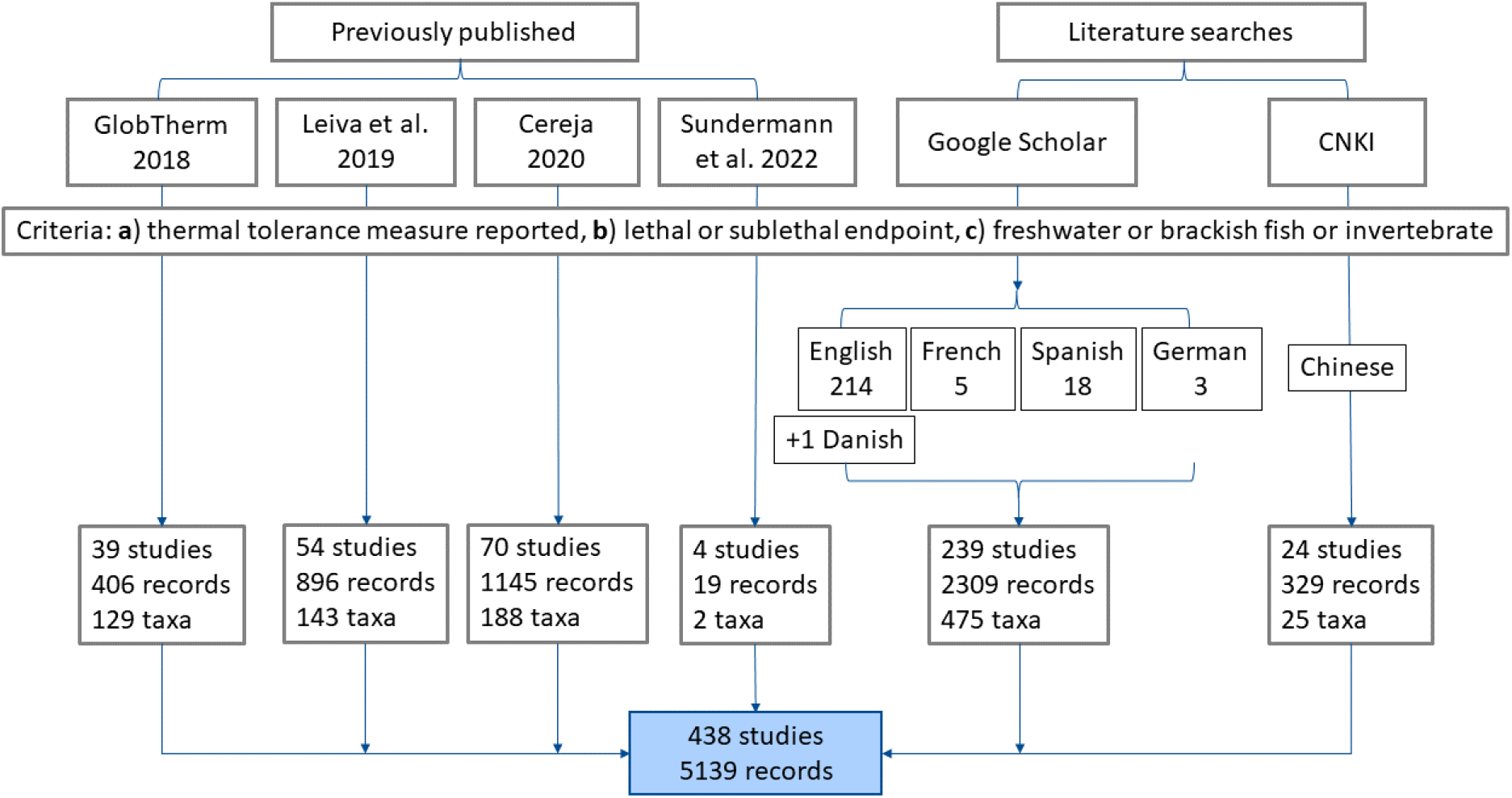
A flow chart describing the workflow to obtain studies included in the database. A total of 438 studies were included which resulted in 5139 records of heat tolerance in total. The numbers below the languages indicate the number of relevant studies for each language. One Danish study which came up in the English language search results was included; 8 additional studies were obtained from references.

### Harmonizing data from existing databases

We selected four recently published databases on thermal tolerance which include freshwater organisms and contain variables relevant to the test organisms and the test metrics. GlobTherm contains thermal tolerance records for the largest number of species in one compilation to date, the compilation of Leiva et al. includes a relatively high number of freshwater invertebrate taxa, the compilation by Cereja includes only aquatic species (therefore more freshwater taxa) and notes additional stressors, and the dataset by Sundermann et al. focuses exclusively on freshwater invertebrates. Other large datasets either contained fewer relevant variables due to the nature of the research questions they were collected for, fewer freshwater taxa (especially invertebrates), or a high duplication of studies with the selected databases.

An initial attempt was made to harmonize the databases selected. However, previous databases either only reported one value per taxon, even when multiple were reported in constituent studies, mislabelled thermal tolerance metrics (i.e. labelling all thermal tolerance values as CTmax, though some were other metrics), or lacked key information related to the method, location, or test organism.

Thus all databases were filtered for relevant freshwater or brackish taxa, all existing data for these combined, and each corresponding source reference examined to cross-check existing data, correct errors, and fill in missing variables. This included extracting additional data from the main text, tables, figures, and supplementary information of source studies. When data was only presented in figures, data was extracted using Plot Digitizer^73^. Collected variables are summarized in **Table 1**, and the extraction protocol with descriptions of all variables in detail is available in the supplementary information (**S1**).

**Table 1.** An overview of the variables included for records in the database.

| Variable type | Variables | Variables queried |
| --- | --- | --- |
| Test organism | Taxon, common name, life stage, sex, body mass, body length, group, habitat, origin | Habitat classification queried from WORMS |
| Taxonomy | Domain, kingdom, phylum, class, order, family, genus, species, subspecies | Queried from the Open Tree of Life |
| Thermal tolerance and method | Tolerance value, metric, sample size, error, error measure, acclimation time, acclimation temperature, ramp rate, test time, endpoint, aeration, pH, salinity, oxygen |  |
| Environmental information | Environmental temperature at collection, environmental | Köppen-Geiger climate region, elevation, continents, and |
|  | temperature maximum/minimum, season, season type, climate region, continent, country, elevation | countries queried from rnaturalearth using coordinates |
| Sampling information | collection year, collection dates, latitude and longitude of collection location |  |
| Stressor information | Stressor type, stressor level, stressor units |  |
| Reference information | Reference code, full reference citation, doi, journal, publication year |  |
| Notes | General notes, metric note, taxon name note, endpoint note, stressor note, body size note, origin note |  |
| Other | Test id, study id, taxon id, location id, ott id | OTT ID queried from the Open Tree of Life using the taxon name |

### Literature searches

Fish made up the majority of freshwater taxa represented in existing databases, with invertebrates vastly underrepresented^43^. The database containing the most taxa (over 2000) includes only 8 species of freshwater insects^43^, a four-fold underrepresentation according to current estimates of the total species on earth^74,75^. Therefore we exerted additional effort in finding papers focused on the thermal tolerance of freshwater invertebrates. Scoping searches confirmed that papers focused on invertebrates were partly missed in searches with general search terms. We modified the general English search terms by adding the order, family, or genus name of freshwater invertebrates (**S2, Table S1**) obtained from a list of freshwater invertebrates sampled in Germany over the last 12 years, spanning habitats from near-natural to highly degraded conditions^76,77^. Though the sampling occurred in only one country in Europe, the orders and families used are distributed globally^78^, and papers resulting from these searches were not limited to one geographic region.

Beside the specific search for literature on freshwater invertebrates, a general literature search was conducted in Google Scholar in English for the time period from January 1^st^ 2019 until April 26^th^ 2023, to cover studies that appeared after the previous databases were compiled. Pilot searches were conducted in all languages to fine-tune the search terms. Final searches in French, German, and Spanish were conducted in Google Scholar, and the search in Chinese was conducted in CNKI, for the earliest indexed studies up until April 26^th^ 2023. The pilot search and search term refinement for the Spanish search was performed in SciElo and Google Scholar, but more relevant publications were indexed in Google Scholar so the final search was conducted there. Final search terms in all languages are recorded in **Table S2** (**S2**). Pilot searches in Italian and Norwegian did not result in any relevant publications. One publication in Danish came up as part of the English search and was included (**Figure 1**). Papers that came up in other languages were excluded.

Google Scholar search results were saved as a file using Publish or Perish software^79^. Results from searches were screened first for relevance by title and abstract, then for duplicates with the studies already included from previous databases. Some studies were eliminated during the data extraction phase because inclusion criteria were not met, in these cases the reason for elimination was noted.

### Data extraction and processing

Data was extracted according to a standardized protocol (**S1**). Following data extraction, a subset of the data was cross-checked to the source references by three authors. Data was also examined for outliers and unreasonable values indicative of errors (for example, a weight of 2 kilogram for invertebrate larvae was reported, which was caused by a missed decimal point) were corrected. Spelling errors were also checked and corrected. The endpoint was scored variably by different authors, though they often were essentially the same. These were unified into simpler categories and details relegated to the notes (**S2, Table S3**). Once datasets from all authors were compiled and errors checked, coordinates were used to query the Köppen-Geiger climate region, elevation, continent, and country of each sampling location. Full taxonomic information for each taxon was queried from the Open Tree of Life^80^, which collates taxonomic information taxonomy from multiple sources. To do this, Open Tree Taxonomy (OTT) identifiers were queried for all taxa names in the database, then taxonomic classifications were added for each taxa using the OTT id. All screening, harmonization, data processing, and querying was conducted in R^81^ with the following packages: revtools^82^, tidyverse^83^, data.table, rotl^84^, taxize^85^, sf^86^, elevatr^87^, geodata^88^, terra^89^, fs^90^ rnaturalearth^91^, viridis^92^.

## Data Records

The database includes 5139 records for a total of 834 species, 452 invertebrates and 382 fish, from 722 locations worldwide (**Figure 2**). Of the 834 total species, 594 reside solely in freshwater, 72 only in brackish waters, and 168 in both. The mean number of tests per species is approximately 6, the median 3, with 606 species having more than one record and 153 more than 10, which allows for assessment of intraspecific variation for a subset of species. A total of 395 tests recorded thermal tolerance in the presence of an additional stressor (chemical, pathogen, reduced oxygen, flow velocity, salinity). The inclusion of variables like body size, life stage, coordinates, and environmental conditions allow for a wide range of hypotheses to be tested with this data. The detailed information on test metrics and methodology allow for metrics to be compared, or for records to be filtered according to desired features, a high or low acclimation temperature, for example. The database features records from 1900 until the present day, which presents the opportunity for comparisons across time, though these may be limited by the scarcity of early data (**Figure 3**). Non-English languages accounted for 12.2% of records, with 6.4% from Chinese, 3.6% Spanish, 0.7% French, 0.4% German, and 0.1% Danish papers. However, the records are not distributed evenly by language across climate regions – Spanish papers contribute 13.6% of arid and 7.8% of tropical climate records, and Chinese 10.7% of continental and 8.5% of temperate climate records within the database (**Figure 2**). Non-English languages added 43 taxa to the database, and additional data for 32 taxa that also had records from English papers; these were from 48 additional locations.

**Figure 2.**
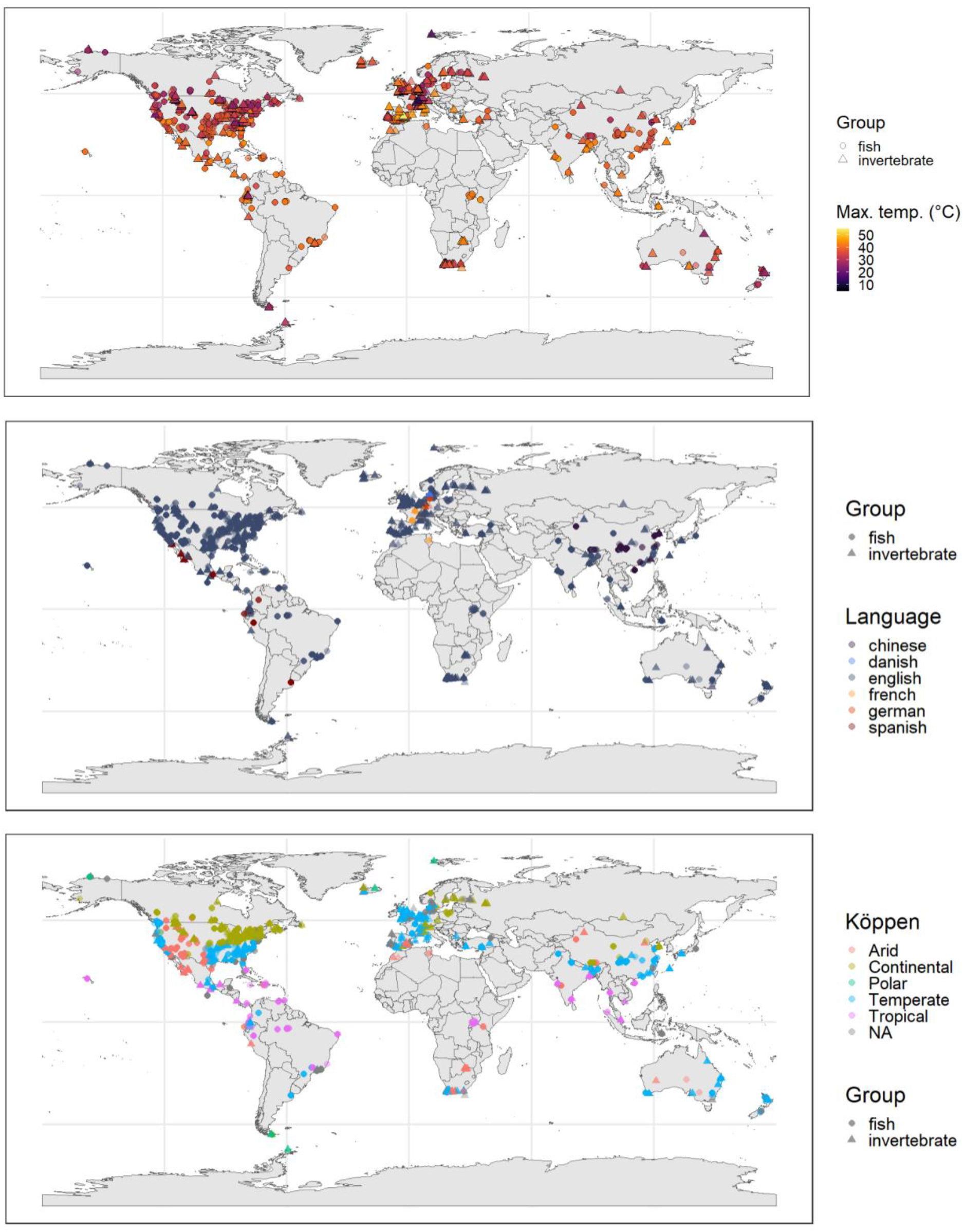
Geographic distribution of test organisms in the database by maximum temperature tolerance, language, and Köppen-Geiger climate classification. A total of 722 locations are represented in the database.

**Figure 3.**
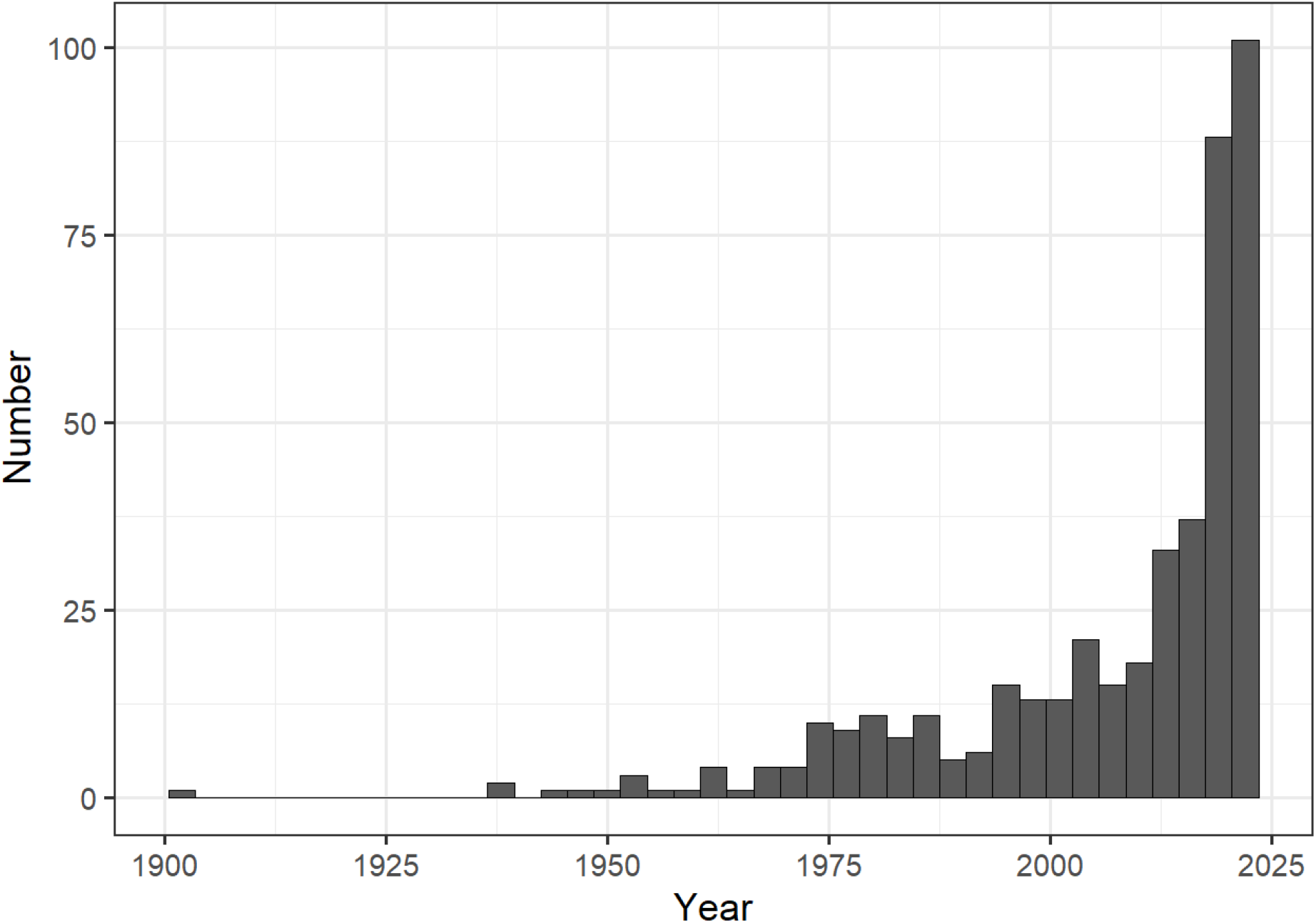
Histogram of the number of studies per publication year included in the database.

## Technical Validation

The concentrated search effort added 295 invertebrate taxa, nearly double the 157 taxa obtained from existing databases. Of a total of 452 freshwater or brackish invertebrates, 313 are insects, representing 0.3% of species estimated to exist^74^. While this may seem low, it is several hundred to thousand-fold more than previous work focused on all taxa^43,48,49^.

A manual cross-check to the original source was done for a subset of the data, with 24.77% of rows cross-checked by at least one person. Additionally the distribution of all numerical variables was visualized in R, and outliers manually checked. Typos in taxon names were corrected, and outdated species names changed to currently accepted names (as of July 2024 in the Open Tree of Life Taxonomy). The names as referred to in the original source are in the notes column where names were changed. The OTT id, which is included as a variable, allows for names to be updated more easily should they change in the future.

All records include a taxon name at the genus or species level, the origin of the test taxon, a thermal tolerance measure, the metric (indicating which methodology was used), the endpoint, the habitat, the location of sampling (or laboratory location for non-wild organisms), variables relating to the location (continent, country, elevation), and reference information (including publication year and language). Over 90% of records include sample size, acclimation information, and life stage of the organism at the time of the test. Error measures are included for 65% of records; the type of error measure is also noted in all cases where error is included. This information, or lack thereof, can be used to filter the data as needed for use in investigating various research questions.

While we attempted to counteract geographic bias in our approach, Europe and North America represent 65.56% of total records in the database though they make up 23.67% of Earth’s land area. With only English studies included, they would make up 69.95% of records. So while non-English languages contribute to lessening data gaps (**Figure 2**), much work remains to be done to fill large gaps in the geographic distribution of data.

When it comes to data gaps, multiple factors, include political ones, are at play. It is certainly possible that data is simply absent from certain areas, due to a lack of access, interest, or resources. In these cases more research and resources are needed in areas that need them most. However, data may also be locked in older studies which are much more difficult to access than more recently published work. For instance, thermal tolerance papers which were concerned with deleterious effects of thermal effluents from power plants are prevalent in the time period from 1960 to 1980, and while scans of these are common in US archives that are indexed by Google Scholar, the archives of other countries, which almost certainly also had research programs on the topic^93^, are difficult to impossible to access for non-native researchers. The research of the largest country in Eurasia, whose land area contains one fifth of the world’s fresh water but also presents a large data gap, is difficult to access not only because of linguistic barriers but also political ones.

## Code Availability

The code to process and aggregate the data, as well as data in raw, processed, and aggregated form, is available on GitHub.

## Acknowledgements

We thank all student helpers who assisted in data extraction. We also thank the entire Collaborative Research Center RESIST for providing the research environment wherein this project developed. We appreciate all authors who authored the original work included, without whom this project could not be completed. Funded by the Deutsche Forschungsgemeinschaft (DFG, German Research Foundation) – SFB 1439/1 2021 – 426547801. FH acknowledges the support from the Chinese Academy of Sciences (E355S122).

## Author contributions

The database was conceptualized by RBS and HSB, with input by FH to make it multi-lingual. HSB developed the search protocol, harmonized existing databases, conducted literature searches in English and German, processed the final dataset, and wrote the manuscript. Data querying from English papers was conducted by HSB, SP, KP, JFJ, JWS, AM and NJK. FH conducted the literature search in Chinese, and extracted the data along with XC. Literature search and extraction for Spanish papers was conducted by GMM and CES. SP conducted pilot searches in Italian and Norwegian, and the search and data extraction in French. HSB, KP, and FH conducted cross-checks of the data extracted by other collaborators. RBS supervised the project and edited the manuscript.

