## Supplement 1 for "Global thermal tolerance of freshwater invertebrates and fish"

**S1. Supplementary information.**

Collecting heat tolerance data protocol

Last updated 03/08/2023

**Description**

Thermal tolerance has been measured for species in various forms since the late 1700s. More recently the metrics for measuring thermal tolerance have been standardized and collected in larger datasets. The figure visualizes the relationship to acclimation temperature (Souchon & Tissot 2012).

The most common methods and metrics for thermal tolerance are:

- ➔ The critical thermal method (CTM), a method where an organism acclimated to a certain temperature (acclimation temperature) is subject to increased heating at a certain rate (the ramp rate) until a physiological endpoint of some sort is observed. The endpoint varies by organism and is not lethal; for example, a common endpoint for fish is loss of equilibrium (LOE) as observed by tipping over/no longer righting correctly in the water. The metrics recorded for this method are the critical thermal maximum (CTmax) for high temperatures and critical thermal minimum (CTmin) for low temperatures. This method is typically conducted multiple times per individual (repeats occur after a recovery period of at least 24 hours).
- ➔ The incipient lethal temperature (ILT) method  is set up similarly to a toxicity test. A pool of organisms acclimated to a certain temperature are split up into groups and put into a container held at a certain constant temperature. A range of temperatures is tested with ~5 organisms per test temperature. Mortality is monitored after a set period of time – 96 hours and 7 days are common test times, but shorter ones are also possible. The lethal temperature of a certain mortality (LTx) is measured – most commonly temperature at which 50% mortality occurred (LT50).

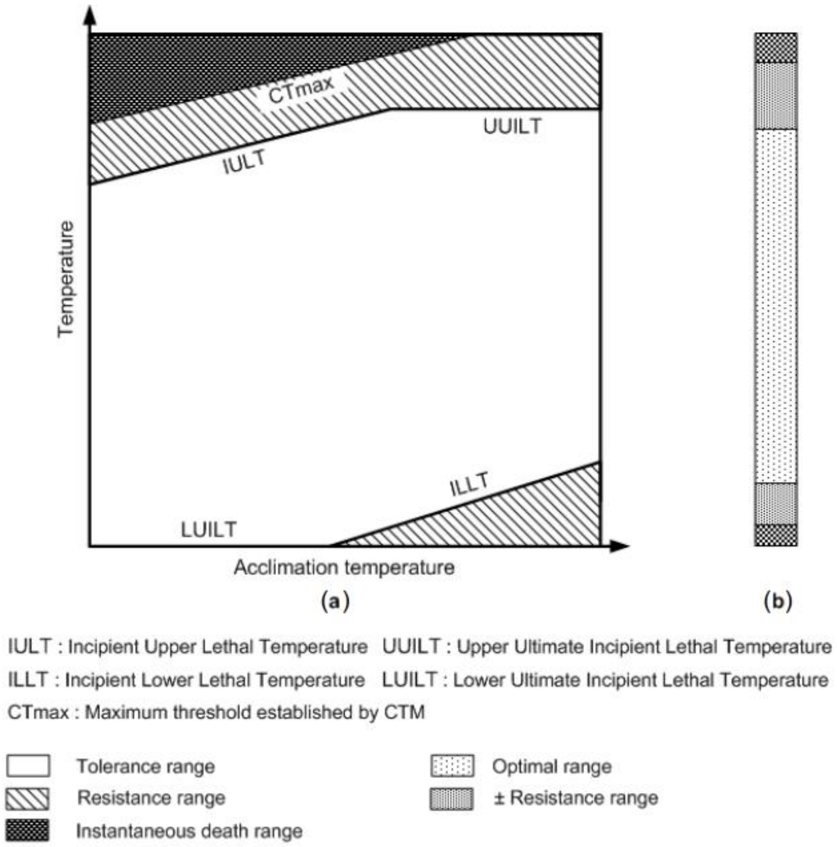


**Aims**

The aim is to collect thermal tolerance values and relevant variables from source studies obtained by systematic literature search.

**Protocol**

The following variables will be collected and compiled from the original source, if available:

| # | Variable name | description |
|---|---|---|
| | person | Name of person querying the data from the source study |
| | taxon | Taxonomic name in Latin (species) |
| | common_name | Common name (if given), in language of reference |
| | taxon_name_notes | Notes on taxon name – separate notes with commas |
| | tmax | The thermal maximum – a temperature in $^{o}$C |
| | n | Sample size for the tolerance value |
| | metric | The metric of the tolerance value – ctmax, lt50 |
| | | If the metric is neither LT50 nor CTmax, write “other” |
| | metric_note | If metric is “other”, write what the authors called it in the source study |
| | error | The error measured on the tolerance value |
| | error measure | The error type measured – standard error (SE), standard deviation (SD), or other metric if neither |
| | acclim temp | The acclimation temperature for the thermal tolerance value – a temperature in $^{o}$C |
| | acclim time | The duration that specimens were kept at the acclimation temperature at before beginning the thermal tolerance tests – in unit recorded in the study |

| | | |
|---|---|---|
| | acclim_time_unit | Recorded unit for acclimation time – d for days, h for hours, min for minutes |
| | start_temp | Starting temperature of the test (only for tests with ramping procedure), most of the time this is the same as acclimation temperature |
| | ramp | The temperature ramp rate for CTmax tests – in $^{o}$C/min |
| | test time | Duration of the test |
| | test_time_unit | Unit of test time/duration as recorded in reference study |
| | endpoint | Which physiological endpoint was measured, mainly relevant for CTmax – loss of movement (LOM), loss of righting reflex (LORR) |
| | temp_tested | For LT50 tests – number of temperatures tested |
| | temp_rep | For LT50 tests – number of replicates per temperature |
| | temp_rep_n | For LT50 tests – number of organisms per replicate |
| | env temp | The temperature of the collection location of the organism – in $^{o}$C |
| | env_temp_sp | Collection type – air, water, point, average – if average, label with daily, weekly, monthly |
| | env_temp_max | Maximum collection temperature if recorded |
| | env_temp_min | Minimum collection temperature if recorded |
| | season | Season during which organisms were collected and thermal tolerance was tested – winter, spring, summer, fall |
| | | if collection and testing occurred in different seasons, enter the season that individuals were collected, unless collected individuals were kept outdoors – then put season during which individuals were tested. For tests where individuals were not wild-collected, enter the season during which the test was conducted.<br>If collected and tested across multiple seasons, put "multiple" |
| | season_type | Can put more than one if collection/testing occurred across multiple seasons (separate with comma) - hot, cold, intermediate |
| | coll start | Put start date of collection, if reported, in ISO format: YYYY-MM-DD<br>If only month is reported, put first day of that month |
| | coll end | Put end date of collection, if reported, in ISO format: YYYY-MM-DD<br>If only month is reported, put last day of that month |
| | coll year | the year during which specimens were collected; if none recorded, leave blank |
| | | If collection occurred over multiple years, put the first one<br>If collection occurred over more than three years and thermal tolerance values are inseparable between collection years, leave blank |
| | origin | Origin of the tested specimens – wild, commercial, or lab |
| | | Eggs collected in the wild and reared in the lab count as wild, but with note "lab_reared", offspring whose mothers were wild collected have origin "lab" with note "first_gen" |
| | origin_note | lab_reared, first_gen |
| | life stage | Life stage of the tested specimens – adult or non-adult |
| | life stage sp | For non-adults: list life stage stated in paper, these can be collated later |
| | sex | male, female, mixed if both explicitly mentioned (doesn't have to be 50:50), blank if unreported |
| | body mass | Body mass of tested specimens – average of the sample in grams |
| | | When range is given, use midpoint |

| | | |
|---|---|---|
| | error bm | Error of the body mass, if reported |
| | error measure bm | Type of error of the body mass, if reported |
| | bm type | Type of body mass measure; wet or dry mass |
| | body length | Body length of tested specimens – average of sample in mm |
| | | When range is given, use midpoint |
| | error bl | Error of the measured body length, if reported |
| | error measure bl | Type of error of the body length, if reported |
| | size notes | Notes on organism size (mass or length) if relevant |
| | climate | Climate of the native range of the species – polar, temperate, subtropical, or tropical |
| | habitat | Habitat of organisms, if recorded – marine, freshwater, brackish |
| | | For fish which switch between marine and freshwater habitats – put water type that they were raised in/collected from |
| | group | Organism group to which the species belongs – fish, invertebrate, algae; exclude amphibians, reptiles, mammals, etc |
| | lat | the latitude of the collection location of the species – in decimal degrees format – use Google Maps for this |
| | long | the longitude of the collection location of the species – in decimal degrees format - use Google Maps for this |
| | | If exact latitude and longitude of the collection location are not available, enter the coordinates for the testing location<br>For non-wild collected specimen, enter coordinates of culture location – for eggs or very young specimens originating from aquaculture/commercial sources that were shipped to the lab and reared there before testing, enter lab location |
| | elevation | The elevation of the collection location of the species – in meters<br>For non-wild caught specimens, enter elevation of lab where tests were conducted |
| | continent | Continent of the collection location of the species – North America, South America, Eurasia, Oceania, Africa, Antarctica |
| | country | Country of collection and testing, if recorded |
| | aeration | If test system is not aerated or oxygen not added, put "No" (for aquatic tests only), if so put "Yes", if water is flowing/circulating but nothing else is mentioned, put C |
| | oxygen | Oxygen concentration |
| | pH | pH of water during test, if recorded |
| | salinity | Salinity of test, if recorded |
| | salinity_unit | Salinity unit if recorded – conductivity or concentration units |
| | add_stressor | indication if tolerance was measured with an additional stressor – put yes or no |
| | type_stressor | type of additional stressor, if present<br>for oxygen put oxygen<br>for chemicals put chemical<br>For parasite infections, put parasite<br>For other infections, put pathogen |
| | stressor_type_sp | Chemical name if chemical stressor present,<br>Parasite name for parasite, pathogen name for pathogen |
| | level_stressor | level of the additional stressor, if present |

| | | |
|---|---|---|
| | | for oxygen, put high/low (not concentration) |
| | level_stressor_unit | Unit of stressor – for chemicals, concentration units |
| | notes_stressor | notes on the additional stressor, if present |
| | plotdigit | Put “T” if data is extracted using plot digitizer |
| | supp | Put “T” if data is found in supplementary information |
| | ref_source | Database that reference was obtained from – GS for Google Scholar, WOS for Web of Science |
| | ref_language | Original language of reference |
| | pub_year | The publication year of the thermal tolerance study |
| | ref | Reference for the thermal tolerance value |
| | | Code as follows:<br>One author: AuthorLastName_Year<br>Two authors: Author1_&_Author2_Year<br>More authors: Author1_et_al_Year<br>For non English studies code this in English |
| | ref info | Full citation of the reference – in English |
| | ref_info_orig | Full citation of the reference – in original language |
| | ref doi | The doi of the reference (should be the same across languages) |
| | ref_journal_orig | Journal of the reference – in original language |
| | notes | Other relevant information |
