## Supplement 2 for "Global thermal tolerance of freshwater invertebrates and fish"

**S2. Supplementary information.**

**Table S1**. List of invertebrate taxonomic names used in additional Google Scholar searches. The respective order, family, and genus names were added to the search string at the top of the table.

| Search string | | |
|---|---|---|
| "**taxonomic name**" AND "thermal tolerance" OR "heat tolerance" OR "ctmax" OR "incipient lethal temperature" OR "critical thermal maximum" OR "thermal limit" | | |
| Order | Family | Genus |
| "Bivalvia" "Coleoptera" "Crustacea" "Diptera" "Ephemeroptera" "Gastropoda" "Heteroptera" "Hirudinea" "Hydrachnidia" "Lepidoptera" "Megaloptera" "Neuroptera/ Planipennia" "Odonata" "Oligochaeta" "Planipennia" "Plecoptera" "Seriata" "Trichoptera" "Turbellaria" | "Aphelocheiridae" "Asellidae" "Astacidae" "Athericidae" "Baetidae" "Beraeidae" "Bithyniidae" "Brachycentridae" "Caenidae" "Calopterygidae" "Cambaridae" "Ceratopogonidae" "Chaoboridae" "Chironomidae" "Chloroperlidae" "Coenagrionidae" "Corbiculidae" "Cordulegastridae" "Corixidae" "Corophiidae" "Culicidae" "Dendrocoelidae" "Dixidae" "Dolichopodidae" "Dryopidae" "Dugesiidae" "Dytiscidae" "Elmidae" "Empididae" "Enchytraeidae" "Ephemerellidae" "Ephemeridae" "Ephydridae" "Erpobdellidae" "Gammaridae" "Gerridae" "Glossiphoniidae" "Glossosomatidae" "Goeridae" "Gomphidae" "Gyrinidae" "Haemopidae" "Haliplidae" "Haplotaxidae" "Helophoridae" "Heptageniidae" "Hydraenidae" "Hydrobiidae" "Hydrometridae" "Hydrophilidae" "Hydropsychidae" "Hydroptilidae" "Janiridae" "Lepidostomatidae" "Leptoceridae" "Leptophlebiidae" "Leuctridae" "Libellulidae" "Limnephilidae" "Limoniidae" | "Agrypnia" "Alboglossiphonia" "Amphinemura" "Amphipoda" "Anabolia" "Ancylus" "Anisus" "Anodonta" "Anomalopterygella" "Antocha" "Aphelocheirus" "Asellus" "Atherix" "Athripsodes" "Atrichopogon" "Atrichops" "Baetis" "Bathyomphalus" "Beraea" "Beraeodes" "Beris" "Bithynia" "Brachycentrus" "Brachyptera" "Brychius" "Caenis" "Calopteryx" "Centroptilum" "Ceraclea" "Ceratopogoninae" "Chaetopterygini" "Chaetopteryx" "Chaoborus" "Chelifera" "Cheumatopsyche" "Chironomini" "Chironomus" "Chrysopilus" "Chrysops" "Clinocerinae" "Cloeon" "Coenagrion" "Corbicula" "Cordulegaster" "Corixinae" "Corophium" "Crunoecia" "Cymatia" "Cyphon" "Cyrnus" "Dasyhelea" "Decapoda" "Dendrocoelum" "Deronectes" "Dicranota" "Dikerogammarus" "Dina" "Dinocras" "Dixa" "Dixella" "Dolichopeza" "Drusus" "Dryops" "Dugesia" "Ecclisopteryx" "Ecdyonurus" "Eiseniella" "Electrogena" "Elmis" "Elodes" |

| | | |
|---|---|---|
| | "Lumbricidae"<br>"Lumbriculidae"<br>"Lymnaeidae"<br>"Micronectinae" "Muscidae"<br>"Nemouridae"<br>"Nepidae" "Notonectidae"<br>"Odontoceridae"<br>"Oligoneuriidae" "Pediciidae"<br>"Perlidae"<br>"Perlodidae"<br>"Philopotamidae"<br>"Phryganeidae" "Physidae"<br>"Piscicolinae" "Planariidae"<br>"Planorbidae"<br>"Platycnemididae"<br>"Polycentropodidae"<br>"Potamanthidae"<br>"Psychodidae"<br>"Psychomyiidae"<br>"Ptychopteridae"<br>"Rhagionidae"<br>"Rhyacophilidae"<br>"Sciomyzidae" "Scirtidae"<br>"Sericostomatidae"<br>"Sialidae" "Simuliidae"<br>"Sisyridae" "Sphaeriidae"<br>"Stratiomyiidae" "Syrphidae"<br>"Tabanidae"<br>"Taeniopterygidae" "Tipulidae"<br>"Unionidae" "Veliidae"<br>"Viviparidae" | "Eloeophila" "Epeorus"<br>"Ephemera" "Ephemerella"<br>"Eristalini" "Erpobdella"<br>"Esolus"<br>"Galba" "Gammarus"<br>"Gerris" "Glossiphonia"<br>"Glossosoma"<br>"Glyphotaelius" "Goera"<br>"Gomphus" "Gyraulus"<br>"Gyrinus" "Habroleptoides"<br>"Habrophlebia" "Haemopis"<br>"Halesus"<br>"Haliplus" "Haplotaxis"<br>"Helius" "Helobdella"<br>"Helochares" "Helophorus"<br>"Hemerodromia"<br>"Hemiclepsis" "Heptagenia"<br>"Hesperocorixa"<br>"Heteroptera" "Hippeutis"<br>"Hydatophylax"<br>"Hydrachnidia"<br>"Hydraena" "Hydrellia"<br>"Hydrocyphon" "Hydrometra"<br>"Hydroporinae" "Hydroporus"<br>"Hydropsyche"<br>"Hydroptila" "Hygrotus"<br>"Ibisia" "Idioptera"<br>"Ilybius" "Ironoquia"<br>"Ischnura"<br>"Isoperla" "Jaera"<br>"Laccobius" "Laccophilus"<br>"Lepidoptera" "Lepidostoma"<br>"Leptocerus"<br>"Leptophlebia" "Leuctra"<br>"Libellula" "Limnebius"<br>"Limnephilini" "Limnephilus"<br>"Limnius"<br>"Limnophora" "Lispe"<br>"Lithax" "Lumbriculus"<br>"Lymnaea" "Lype"<br>"Micrasema"<br>"Micronectinae" "Micropterna"<br>"Molophilus" "Musculium"<br>"Mysidacea" "Mystacides"<br>"Nebrioporus"<br>"Nemoura" "Nemurella"<br>"Nepa" "Nephrotoma"<br>"Odontocerum" "Oecetis"<br>"Oecismus"<br>"Oligochaeta"<br>"Oligoneuriella"<br>"Ophiogomphus"<br>"Orconectes" "Orectochilus"<br>"Oreodytes" "Orthetrum"<br>"Oulimnius"<br>"Paraleptophlebia" "Pedicia" |

| | | |
|---|---|---|
| | | "Pericoma" "Perlodes" "Philopotamus" "Phylidorea" "Physella" "Pilaria" "Piscicola" "Pisidium" "Planorbis" "Platambus" "Platycnemis" "Plecoptera" "Plectrocnemia" "Polycelis" "Polycentropus" "Pomatinus" "Potamanthus" "Potamophylax" "Potamopyrgus" "Proasellus" "Procloeon" "Prodiamesa" "Prosimulium" "Protonemura" "Pseudolimnophila" "Psychomyia" "Ptychoptera" "Pyrrhosoma" "Radix" "Rhantus" "Rhithrogena" "Rhyacophila" Rhypholophus" "Riolus" "Sericostoma" "Serratella" "Sialis" "Silo" "Simulium" "Siphonoperla" "Sisyra" "Sphaerium" "Stagnicola" "Stenophylax" "Stylodrilus" "Tanypodinae" "Tanytarsini" "Tetanocera" "Theromyzon" "Tinodes" "Tipula" "Torleya" "Trocheta" "Turbellaria" "Unio" "Velia" "Viviparus" "Wormaldia" |

**Table S2**. Search terms used in literature searches for each language and platform.

| Language | Search string | Group | Search platform | Conducted by |
|---|---|---|---|---|
| Chinese | ('温度临界值' + '临界温度' + '临界高温' + '临界低温' + '耐热' + '耐冷' + '耐高温' + '耐寒' + '温度耐受' + '高温致死' + '低温致死' + '温度致死')*('鱼' + '鲟' + '鳗' + '鲤' + '鲱' + '鲑' + '鲇' + '鲈' + '鳉' + '鲻' + '鲈' + '鲀' + '鳅' + '鲂' + '鳊' + '鲚' + '鳘' + '鲌' + '鱎' + '鲴' + '鲃' + '鰔' + '鲮' + '鮈' + '鲫' + '鱲' + '鲥' + '鱵' + '鳟' +'鳤' + '鯮' + '鳡' + '鮀' + '鳑鲏' + '鱊' + '鮹' + '鮈' + '鳈' + '鲿' + '鳠' + '䱅' + '鮡' + '鰋' + '鳜' + '鳢' + '鳎') | Fish | CNKI | FH |
| Chinese | ('温度临界值'+'临界温度'+'临界高温'+'临界低温'+'耐热'+'耐冷'+'耐高温'+'耐寒'+'温度耐受'+'高温致死'+'低温致死'+'温度致死')*('无脊椎'+'底栖'+'水生昆虫'+'贝'+'蚌'+'蛤'+'螺'+'腹足'+'瓣鳃'+'虾'+ '蟹'+'蜉蝣'+ '蜉'+'襀翅目' +'石蝇'+'毛翅目'+'石蛾'+'双翅目'+'摇蚊'+'蜻蜓'+'涡虫'+'蚓'+'蛭'+'水龟甲'+'泥甲'+'稻象'+'水象'+'龙虱'+ '溪'+'河'+'湖'+'湿地'+'池塘'+'海'+'淡水') | Invertebrates | CNKI | FH |
| English | ‘ctmax’ AND ‘freshwater’ AND ‘aquatic’ AND ‘critical thermal maximum’ | Fish and invertebrates | Google Scholar | HSB |
| French | "poisson" AND "eau douce" AND "tolérance thermique" OR "tolérance aux chaleur" OR "ctmax" OR "température létale" OR "maximum thermique" OR "limite thermique" | Fish | Google Scholar | SP |
| French | "invertébré" AND "eau douce" AND "tolérance thermique" OR "tolérance à la chaleur" OR "ctmax" OR "température létale" OR | Invertebrates | Google Scholar | SP |

| | | | | |
|---|---|---|---|---|
| | "maximum thermique" OR "limite thermique" | | | |
| German | physiologie tier AND temperaturmaxima OR hitzetoleranz OR hitzeresistenz OR letaltemperatur OR temperaturlimit | Fish and invertebrates | Google Scholar | HSB |
| Spanish | "TCmax" OR "CTMax" OR "Tolerancia+termica" OR "Temperatura+letal" OR "fisiologia+termica" AND "peces" OR "Pez" OR "invertebrados" OR "crustacea" OR "insecta" OR "mollusca" OR "camarón” | Fish and invertebrates | Google Scholar | GMM, CES |

**Table S3**. The endpoint scored during data extraction and the unified endpoint for the final database. All details were added to the endpoint note, and endpoints that were manually checked in the source reference method are labelled.

| endpoint recorded | n | unified | manually checked |
|---|---|---|---|
| 100% mortality | 3 | mortality | yes |
| Absence of righting response | 6 | LRR | |
| Activity decrease | 6 | other | yes |
| Activity increase | 6 | agitation | |
| Cessation of operculum pulsation | 6 | mortality | yes |
| Comma state | 6 | LOM | |
| Complete Equilibrium loss | 22 | LOE | |
| Curling | 12 | LOE | yes |
| DT (desorientacion total) | 42 | LOE | |
| Death | 433 | mortality | |
| Equilibrium loss | 77 | LOE | |
| FO | 32 | OS | yes |
| First equilibrium loss | 6 | LOE | |
| Fish is incapable of holding up | 8 | LOE | |
| Foot detachment | 4 | LOE | |
| Increased locomotion | 1 | agitation | |
| L0RR | 20 | LRR | |
| LOE | 3246 | LOE | |
| LOM | 341 | LOM | |
| LORR | 173 | LRR | |
| LRR | 195 | LRR | |
| Lack of response to tactile stimulus | 5 | LOM | |
| Lethal Temperature | 2 | mortality | |
| Lethal temperature | 5 | mortality | |
| Loss of Orientation | 12 | LOE | |
| Loss of attachment | 5 | LOE | |
| Loss of equilibrium | 164 | LOE | |
| Loss of equilibruim | 6 | LOE | |
| Loss of mobility | 19 | LOM | |
| Loss of movement | 4 | LOM | |
| Loss of operculum movement | 4 | mortality | |
| Loss of responsivness | 3 | LOM | |
| Loss of swimming capacity | 6 | LOE | yes |
| Mortality | 5 | mortality | |
| Muscular spasms | 6 | OS | |
| No heartbeat | 2 | mortality | |
| No movement | 2 | LOM | |
| OS | 95 | OS | |
| Only movement is from operculum | 8 | other | yes |
| Onset of tetany | 8 | LOM | |

| | | | |
|---|---|---|---|
| Response to touch | 2 | LOM | |
| Sporadic fast activity and muscular spasms | 6 | OS | |
| Stop swimming | 36 | LOE | yes |
| Survival for 10 minutes | 12 | survival | yes |
| Survival for 7 days | 12 | survival | yes |
| Total disorientation | 6 | LOE | |
| Valve closure | 8 | mortality | yes |
| agitation | 2 | agitation | |
| collapse of the animal when rolled over onto their sides | 8 | LOE | |
| coma | 24 | LOM | |
| death | 99 | mortality | |
| decrease in pleopod beats | 18 | LOE | yes |
| first retraction into shell | 6 | LOM | yes |
| fish loses the hability to keep a normal position | 8 | LOE | |
| foot curl, loss of grip | 2 | LOE | |
| gaping response | 12 | LOE | yes |
| heat coma | 3 | LOM | |
| inactivity | 32 | LOM | yes |
| irregular undulation | 24 | LOE | yes |
| lack of response | 58 | LOM | |
| lack of response to stimuli | 13 | LOM | |
| lack of response to touch, loss of grip | 2 | LOM | |
| loss in motor coordination | 3 | LOE | |
| loss of attachment | 7 | LOE | |
| loss of coordinated muscle function | 24 | LOE | |
| loss of gaping | 4 | LOM | yes |
| loss of gill movement | 6 | LOM | yes |
| loss of grip | 75 | LOE | |
| loss of movement | 49 | LOM | |
| loss of response | 8 | LOM | |
| loss of righting | 8 | LOE | |
| loss of tube feet activity, righting ability | 2 | LOE | |
| mortality | 822 | mortality | |
| mortality (cessation of pleopod movement) | 42 | mortality | |
| mortality (heart rate decline) | 4 | mortality | |
| onset of muscular spasms | 11 | OS | |
| pleopod movement stopped | 4 | mortality | |
| rapid drop in heart beat | 4 | mortality | yes |
| reproduction | 1 | other | |
| shell emersion | 4 | LOE | yes |
| survival | 8 | survival | yes |
| sustained aerobic swimming | 2 | other | yes |
| total disorientation | 4 | LOE | |
